# Overcoming Protein A-driven Nonspecific Antibody Staining of *S. aureus* in Immunofluorescence Microscopy

**DOI:** 10.64898/2026.03.26.713373

**Authors:** Lena Gauthier, Bettina Löffler, Marc Thilo Figge, Christina Ehrhardt, Christian Eggeling

## Abstract

The ability to detect host cell factors during *Staphylococcus aureus* infection *in vitro* by immunofluorescence microscopy is severely hampered by *staphylococcal* protein A (SpA), a cell wall-anchored protein that binds the fragment crystallizable (Fc) region of immunoglobulins. This interaction generates strong nonspecific fluorescent signals on the bacterial surface, complicating data interpretation and limiting the accuracy of quantitative image analysis. Several measures have been put forward to overcome this obstacle, most importantly the pre-incubation with an anti-SpA antibody (αSpA) and the use of human serum (HS) as blocking agent and antibody diluent. To highlight this feature to general fluorescence microscopy users, we here systematically evaluated these two strategies. Using *S. aureus* coated on coverslips and *S. aureus*-infected A549 cells, we highlight the efficiencies of both approaches to markedly reduce nonspecific fluorescence, with HS treatment yielding the most profound suppression. Notably, HS, containing high levels of human immunoglobulins, offered a robust, cost-effective and broadly applicable solution for minimizing SpA-driven artifacts, thereby improving immunofluorescence microscopy in *S. aureus* infection models *in vitro*.

## Introduction

*Staphylococcus aureus* is a Gram-positive bacterium that asymptomatically colonizes approximately 30% of healthy individuals, primarily residing in nasal passages and on skin.^[1, 2]^ Despite its commensal nature, *S. aureus* is an opportunistic pathogen capable of causing a wide spectrum of diseases, ranging from minor skin and soft-tissue infections to severe systemic illnesses, including pneumonia, sepsis, and endocarditis.^[3, 4]^ In the industrialized world, annual incidence rates of *S. aureus* bacteremia range from 10 to 50 per 100 000,^[4, 5]^ while, globally, over 1 000 000 deaths have been attributed to *S. aureus* in 2019,^[6]^ underscoring its clinical significance and public health impact.

The ability to transition from a harmless colonizer to an invasive pathogen is driven by a complex interplay of host susceptibility, environmental factors, and a diverse repertoire of virulence determinants including immune-modulatory proteins, toxins, and adhesins.^[7-9]^ Among these, *staphylococcal* protein A (SpA)^[10, 11]^ is a key immune-evasive factor that is anchored to the bacterial cell wall^[12-14]^ but also released into the extracellular environment during bacterial growth.^[15]^ SpA is particularly effective due to its dual immunoglobulin-binding activities. On the one hand, released SpA binds to the fragment antigen-binding (Fab) region of variable heavy 3 (VH3) idiotype antibodies, including IgM, IgG, IgA, and IgE,^[16-24]^ thereby disrupting the adaptive immune response.^[25-30]^ On the other hand, cell wall-anchored SpA interferes with the innate immune system by engaging the fragment crystallizable (Fc) region of IgG1, IgG2, and IgG4 from several mammalian species,^[31-39]^ impairing opsonization and Fc receptor-mediated phagocytosis.^[40-43]^ This latter characteristic presents a significant obstacle in laboratory studies utilizing antibody-based detection methods, including ELISA,^[44]^ flow cytometry,^[45]^ or immunofluorescence microscopy.^[46, 47]^

Immunofluorescence (IF) microscopy is a valuable tool, enabling spatially resolved analysis of pathogen-infected cells by labeling defined target molecules with corresponding primary and fluorescently tagged secondary antibodies. However, nonspecific antibody interactions can introduce unintended artifacts. In particular, in the context of *S. aureus* infections, binding of surface-localized SpA to the antibody Fc region generates substantial nonspecific signals,^[46-48]^ confounding accurate quantification of bacterial and host cell components. Therefore, reliable analysis requires strategies to prevent SpA-mediated interference.

Although this issue is well recognized within the *S. aureus* and immunology research communities, and several technical solutions have been proposed, this problem may be less familiar to researchers from different fields, such as biophysics or microscopists. Moreover, comparatively few studies have systematically addressed strategies to overcome SpA-mediated interference in fluorescence-based imaging approaches. For example, Tan *et al*. employed purified human IgG to mitigate SpA-mediated nonspecific staining of *S. aureus*. While the blocking efficacy was quantified for fixed *S. aureus* on microscope slides, blocking in patient tissue sections was only qualitatively assessed.^[47]^ In a separate study, Flannagan and Heinrichs used human serum (HS) as blocking agent and antibody diluent in *S. aureus*-infected cells *in vitro*, though without quantitative evaluation of blocking efficacy.^[49]^

To our knowledge, no previous study has systematically quantified and directly compared the performance of different SpA-blocking strategies in IF microscopy of *S. aureus*-infected cells. As a consequence, comprehensive and accessible methodological guidance – particularly for investigators outside the immediate *S. aureus* field –remains limited, underscoring the need for optimized protocols and broader dissemination of effective solutions.

In this work, we rigorously examined the efficacy of pre-incubation with HS or αSpA under defined *in vitro* conditions. We first optimized the pre-incubation and staining protocols using *S. aureus* coated on coverslips, enabling precise evaluation of blocking efficacies in a simplified system. We then applied the optimized protocols to *S. aureus*-infected cells, demonstrating their applicability in a biologically relevant context. Importantly, we provide detailed experimental procedures in the supporting information to facilitate adoption and reproducibility across laboratories.

Together, our findings provide a robust and reproducible strategy for overcoming a key technical limitation in the imaging of *S. aureus*-infected samples. By substantially reducing nonspecific fluorescence signals, these methods not only improve qualitative visualization but also enable quantitative immunofluorescence analyses, thereby enhancing the utility of antibody-based imaging approaches for studying *S. aureus* pathogenesis and host-pathogen interactions.

## Results

### SpA-mediated Nonspecific Antibody Binding to *S. aureus* Impairs Immunofluorescence Microscopy

To illustrate the challenge posed by the Fc binding activity of SpA during immunofluorescence microscopy, we performed super-resolution stimulated emission depletion (STED) microscopy on *S. aureus* samples (Figure 1). Green fluorescent protein (GFP)-expressing *S. aureus* was stained with an unrelated primary antibody targeting a viral protein (nucleoprotein, NP), which is not expected to bind to the bacterial surface, and secondary antibodies conjugated to Alexa Fluor 568 (AF568) or Alexa Fluor 647 (AF647). The resulting staining pattern was compared to that obtained using a primary antibody specifically targeting SpA. Notably, the unrelated primary anti-NP antibody (αNP) produced substantial nonspecific labeling of the *S. aureus* surface, irrespective of the secondary antibody used, and this nonspecific fluorescence was indistinguishable from the pattern observed when using αSpA as primary antibody (Figure 1).

**Figure 1:**
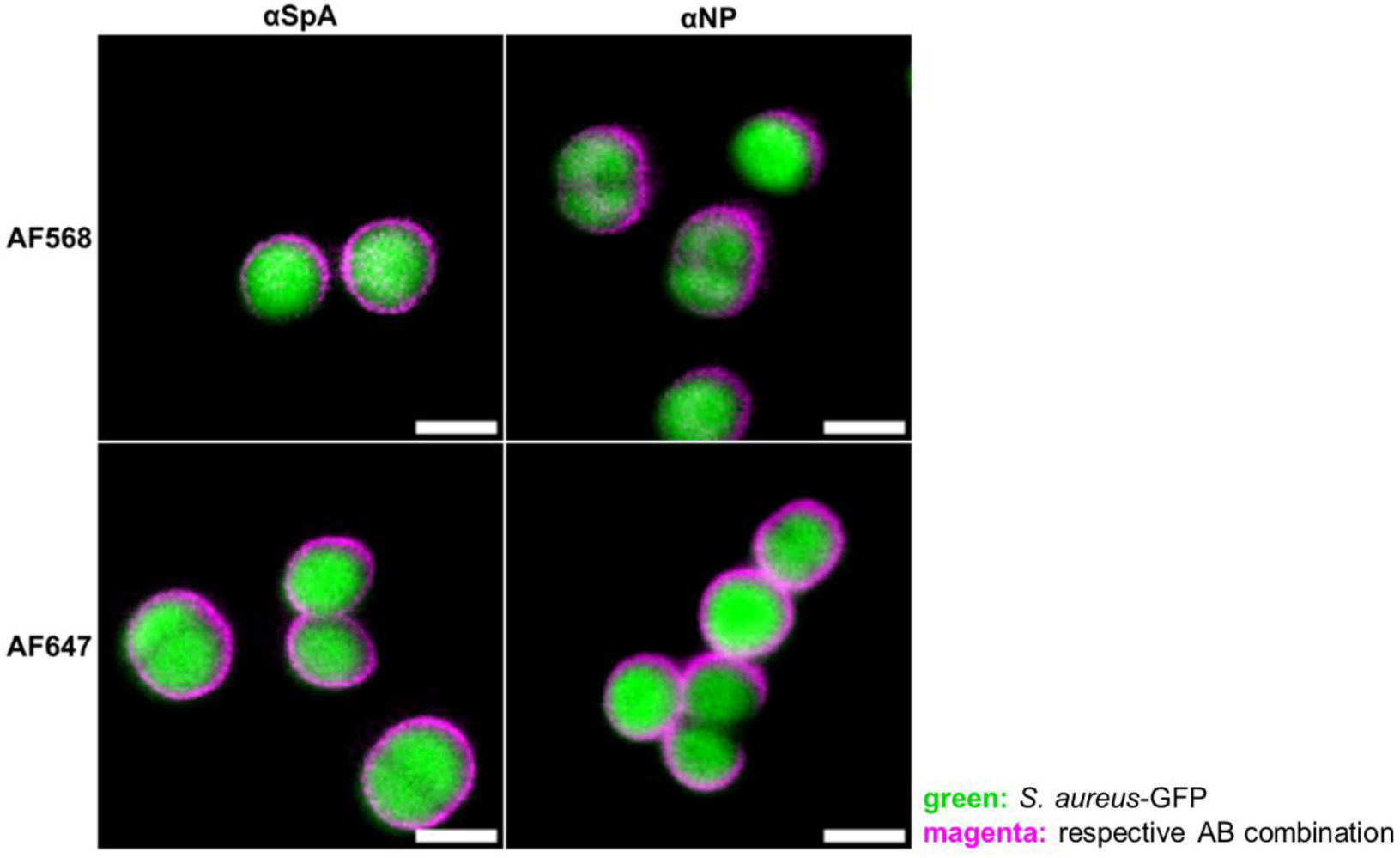
Comparison of SpA-specific and nonspecific antibody staining of *S. aureus*. Representative super-resolution STED microscopy images of GFP-expressing *S. aureus*. Coverslip-coated *S. aureus* was stained with primary antibodies targeting the bacterial surface protein SpA (αSpA, left column) or the unrelated viral protein NP (αNP, right column), and secondary antibodies conjugated with AF568 (top row) or AF647 (bottom row). Scale bars = 1 µm; AB: antibody.

To further demonstrate the generality of this phenomenon, we tested additional unrelated primary antibodies targeting mammalian and viral antigens in a wide field fluorescence microscopy setup. In parallel, we analyzed bacteria stained only with secondary antibodies, and control samples lacking both primary and secondary antibodies. We observed nonspecific fluorescence decorating the bacterial surface for all antibody combinations tested and even in the absence of primary antibodies, while no surface labeling was detected in control samples without antibodies (Supplemental Figure S1). This pervasive nonspecific staining is attributed to SpA’s Fc binding ability, regardless of the antibody target specificity. Thus, even when specific antibodies do not recognize any bacterial epitope, the surface-localized SpA captures all antibodies through its Fc binding activity, resulting in apparent but misleading fluorescence.

### Quantification of the Nonspecific Antibody Staining of *S. aureus*

To systematically evaluate methods aimed at mitigating nonspecific antibody staining of *S. aureus*, it is essential to establish a reliable quantitative readout to assess both the extent of nonspecific labeling and the effectiveness of blocking strategies. To this end, we developed two image analysis pipelines for fluorescent wide-field images to measure the nonspecific fluorescence signals in GFP-expressing *S. aureus* coated on coverslips (Figure 2A-B) and in *S. aureus*-infected cells (Figure 2C-D). Briefly, fixed samples were stained with the unrelated primary αNP and a secondary AF647-conjugated antibody, with Hoechst 33342 counterstaining in the case of *S. aureus*-infected cells. Images were acquired as z-stacks, deconvolved, and processed for quantification. For *S. aureus*-coated on coverslips, maximum intensity projections (MIPs), which clearly depict individual bacterial cocci, were employed (Figure 2A). Bacterial regions of interest (ROIs) were segmented from the GFP-channel, where each ROI corresponds to a single coccus. These ROIs were then overlaid onto the red channel containing the nonspecific fluorescence signal derived from antibody labeling, to measure the mean red intensity per bacterial ROI (Supplemental Figure S2, steps 1-9). Quantification results revealed high levels of nonspecific fluorescence ranging from 10^3^ to 10^4^ a.u. across all ROIs while control samples without any antibodies showed only weak background signals in the range of 10 to 20 a.u. (Figure 2B). Additional quantification of the nonspecific fluorescence observed for the various antibody combinations shown in Supplemental Figure S1 had similar results. When both primary and secondary antibodies were applied, the measured nonspecific signals ranged from 10^2^ to 10^4^ a.u., depending on the respective antibody combination (Supplemental Figure S3A-B). For samples only stained with secondary antibodies, the detected nonspecific signals were subjected to a greater range of 10^1^ to 10^4^ a.u. (Supplemental Figure S3C-D).

**Figure 2:**
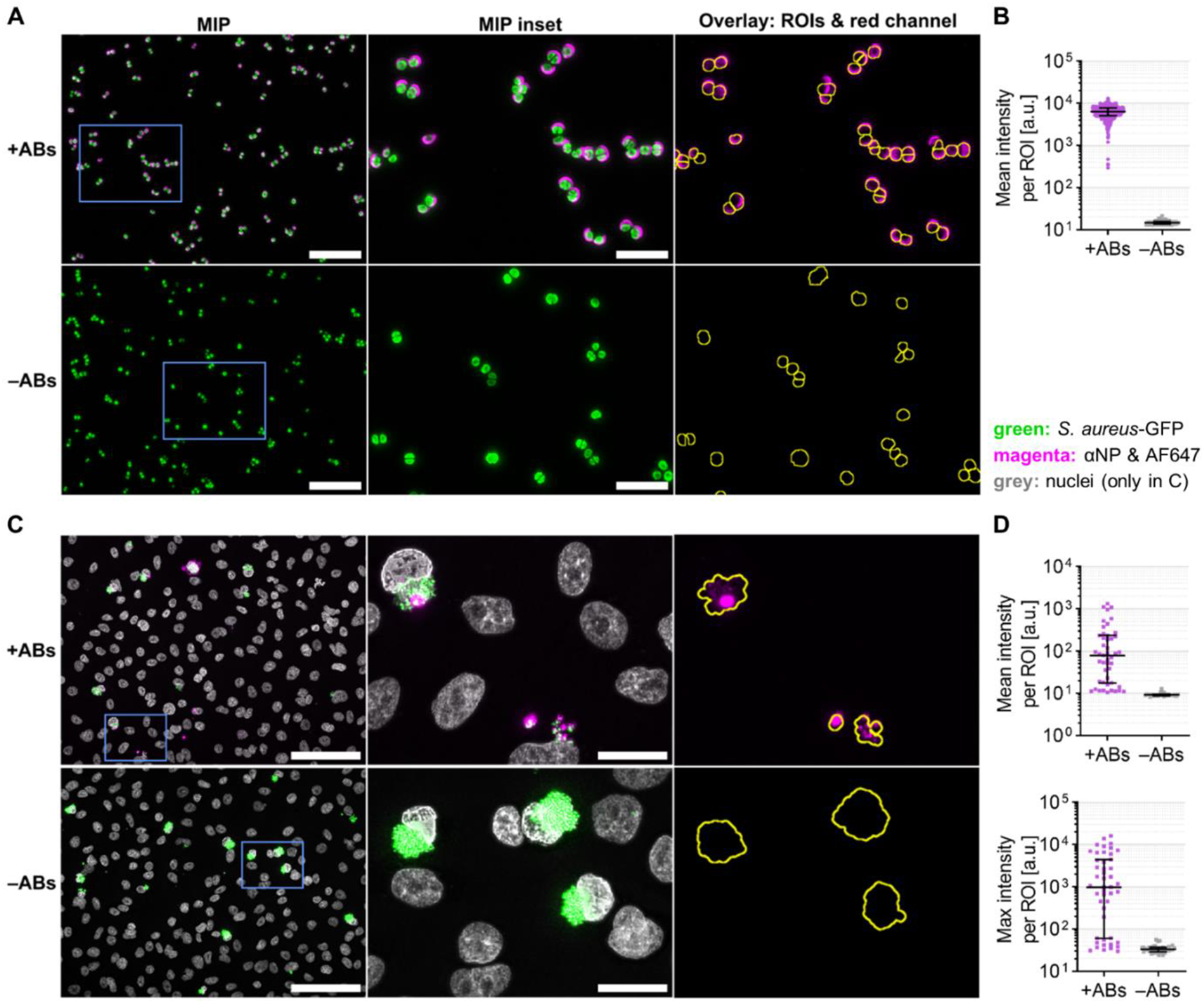
Quantification of the nonspecific antibody staining of *S. aureus*. (A-B) Coverslip-coated GFP-expressing *S. aureus* was stained with αNP and a secondary AF647-conjugated antibody (+ABs) or without antibodies (–ABs). Scale bars = 10 µm (MIPs) or 5 µm (MIP insets). (C-D) A549 cells were infected with GFP-expressing *S. aureus* and stained as in (A) with additional Hoechst 33342 counterstaining. Scale bars = 100 µm (MIPs) or 20 µm (MIP insets). (A, C) Representative wide-field microscopy images. Of note, (C) shows MIPs for better contrast but segmentation and quantification were done in 3D. (B, D) Quantification of the nonspecific signal as described in Figures S2 (B) and S4 (D). Five images per condition were analyzed. Line and error bars depict median ± interquartile range. (D) The two graphs show the mean (top) and maximum (bottom) red intensity per ROI as two alternative read-outs of the same data set. a.u.: arbitrary units; ABs: antibodies; max: maximum; MIP: maximum intensity projection; ROI: region of interest.

In infected cells, segmentation and quantification were performed in all three spatial dimensions (3D) to account for the clustering of *S. aureus* after 9.5 h of infection (Supplemental Figure S4, steps 1-13). In contrast to *S. aureus* coated on coverslips, only parts of the intracellular *S. aureus* clusters exhibited nonspecific staining (Figure 2C), potentially due to biofilm like aggregation involving SpA’s Fc binding activity,^[50]^ which may physically impede antibody access. Consequently, the measured mean red fluorescence intensities per bacterial ROI showed greater variability than those measured for individual cocci on coverslips. To minimize the impact of variable cluster sizes on the quantification, we opted to use the maximum rather than the mean fluorescence intensity per ROI as alternative readout (Figure 2D, bottom panel). Using this approach, clusters with nonspecific staining displayed maximum red intensities ranging from 10^2^ to 10^4^ a.u., while some clusters remained entirely unstained, exhibiting values between 20 and 50 a.u., comparable to the background levels observed in no-antibody controls.

### Pre-incubation with αSpA Blocks Nonspecific Antibody Staining of *S. aureus*

To prevent SpA-mediated nonspecific antibody binding to *S. aureus*, we evaluated two approaches designed to block the Fc-binding sites of SpA prior to the introduction of primary antibodies targeting non-bacterial structures of interest. Our initial approach involved pre-incubating fixed *S. aureus* samples with αSpA before applying the primary and secondary antibodies of interest.

In a first test, fixed *S. aureus* coated on coverslips was pre-incubated with increasing αSpA concentrations prior to staining with the unrelated αNP/AF647 antibody combination. Wide-field fluorescence images revealed a dose-dependent reduction of nonspecific red signals with increasing αSpA concentrations (Figure 3A) and quantitative analysis confirmed this observation (Figure 3B). Without αSpA pre-incubation, nonspecific fluorescence in the range of 10^3^ to 10^4^ a.u. per ROI was detected, while a marked decrease down to the 10^1^ to 10^2^ a.u. range was observed when pre-incubating with a 1:100 or 1:50 αSpA dilution (Figure 3B, Supplemental Figure S5).

**Figure 3:**
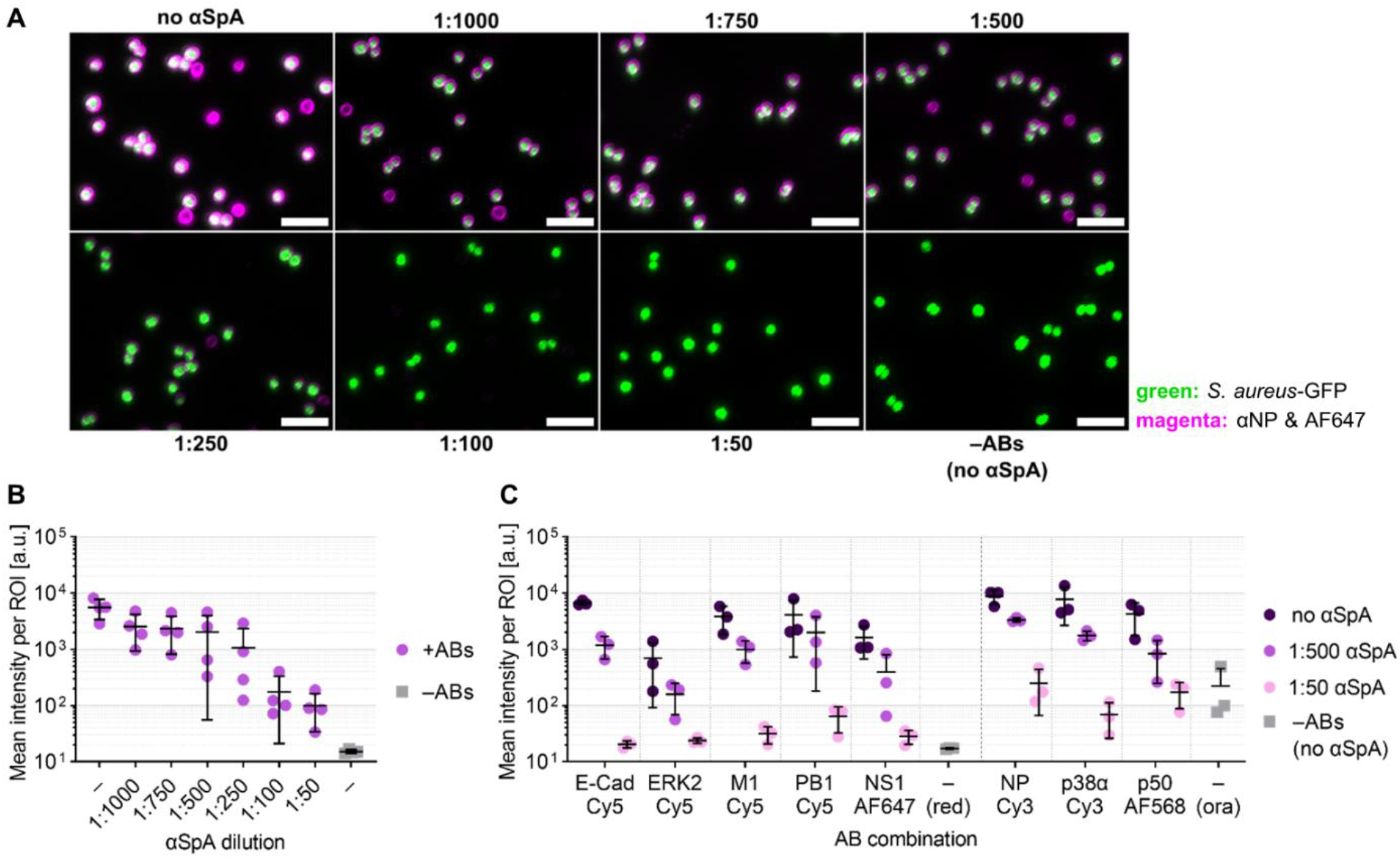
Pre-incubation with αSpA blocks nonspecific antibody staining of *S. aureus*. Coverslip-coated GFP-expressing *S. aureus* was pre-incubated with different αSpA dilutions (1:1000 to 1:50) or with 3% BSA (no αSpA or –) and stained with primary and secondary antibodies (+ABs), or without antibodies (–ABs). (A) Representative wide-field microscopy images. Scale bars = 5 µm. (B-C) Quantification of nonspecific fluorescence as described in Supplemental Figure S2. Nonspecific signals derived from αNP/AF647 staining (B) or from various primary/secondary antibody combinations (C). Five images per condition were analyzed for each biological replicate. Depicted data points show medians of four (B; Supplemental Figure S5) or three (C; Supplemental Figure S6) biological replicates; line and error bars show overall mean ± standard deviation. a.u.: arbitrary units; ABs: antibodies; E-Cad: E-Cadherin; ERK2: extracellular signal-regulated kinase 2; M1: matrix protein 1; MIP: maximum intensity projection; NP: nucleoprotein NS1: non-structural protein 1; ora: orange; p38α: Mitogen-activated protein kinase 14; p50: p50 subunit of NFκB; PB1: polymerase basic protein 1; ROI: region of interest; SpA: *staphylococcal* protein A.

Given the efficacy of this blocking approach for the αNP/AF647 combination, we sought to assess whether the blocking effect was generalizable to other antibody combinations. Specifically, we measured the nonspecific fluorescence signals for several unrelated primary antibodies combined with different secondary antibodies conjugated to red (Cy5 or AF647) or orange (Cy3 or AF568) chromophores. Quantification revealed that pre-incubation with αSpA effectively reduced the nonspecific labeling across all tested combinations (Figure 3C, Supplemental Figure S6).

However, despite the significant reduction in nonspecific fluorescence achieved with αSpA pre-incubation, we observed residual fluorescence between 30 to 200 a.u. per ROI even at the highest αSpA concentration tested (1:50 dilution), which is more than twice as high as the minimal background of 10 to 20 a.u. per ROI seen in no-antibody controls. This prompted us to explore whether an alternative approach could yield even better blocking efficacies.

### HS is a Potent Blocking Agent and Antibody Diluent to Prevent Nonspecific Antibody Staining of *S. aureus*

As highlighted, although pre-incubation with αSpA significantly reduced SpA-mediated nonspecific antibody staining of *S. aureus*, residual fluorescence signals remained detectable. To further enhance the blocking efficacy, we evaluated whether pre-incubation with HS, which contains milligram amounts of IgG per milliliter,^[51-53]^ could achieve superior suppression of nonspecific fluorescence. To systematically assess the impact of pre-incubation with HS, we examined different pre-incubation durations ranging from 1 to 3 h alongside with different antibody diluents (Figure 4). In the previous approaches (Figure 1 to Figure 3), antibodies were always diluted in 3% bovine serum albumin (BSA) but here we additionally tested 50% HS, similar to work by Flannagan and Heinrichs who used human plasma for blocking and antibody dilution when labeling *S. aureus*-infected macrophages with antibodies.^[49]^ Additionally, we considered whether human serum albumin (HSA) had similar effects. Wide-field microscopy images indicated a marked reduction of nonspecific fluorescence across all tested incubation times upon pre-incubation with HS, with minimal differences observed between 1 to 3 h (Figure 4A). However, quantitative analysis revealed residual nonspecific signals in the range of 30 to 80 a.u. per ROI with antibodies diluted in BSA or HSA (Figure 4B). This closely resembled those results obtained with the 1:50 αSpA dilution in our previous approach, where antibodies were always diluted in 3% BSA (Figure 3). Strikingly, the most profound suppression of nonspecific fluorescence down to 10 to 20 a.u. per ROI was achieved when antibodies were diluted in 50% HS, even in the absence of a dedicated pre-incubation step (Figure 4B). Under these conditions, the nonspecific signal was reduced to fluorescence levels comparable to no-antibody controls (10 to 20 a.u. per ROI). These findings highlight a potent blocking capacity of HS and indicate that antibody dilution in IgG-rich HS alone is sufficient to effectively outcompete SpA-mediated nonspecific antibody binding to *S. aureus*. Thus, HS provides a simple yet highly efficient strategy to eliminate SpA-mediated interference in immunofluorescence microscopy.

**Figure 4:**
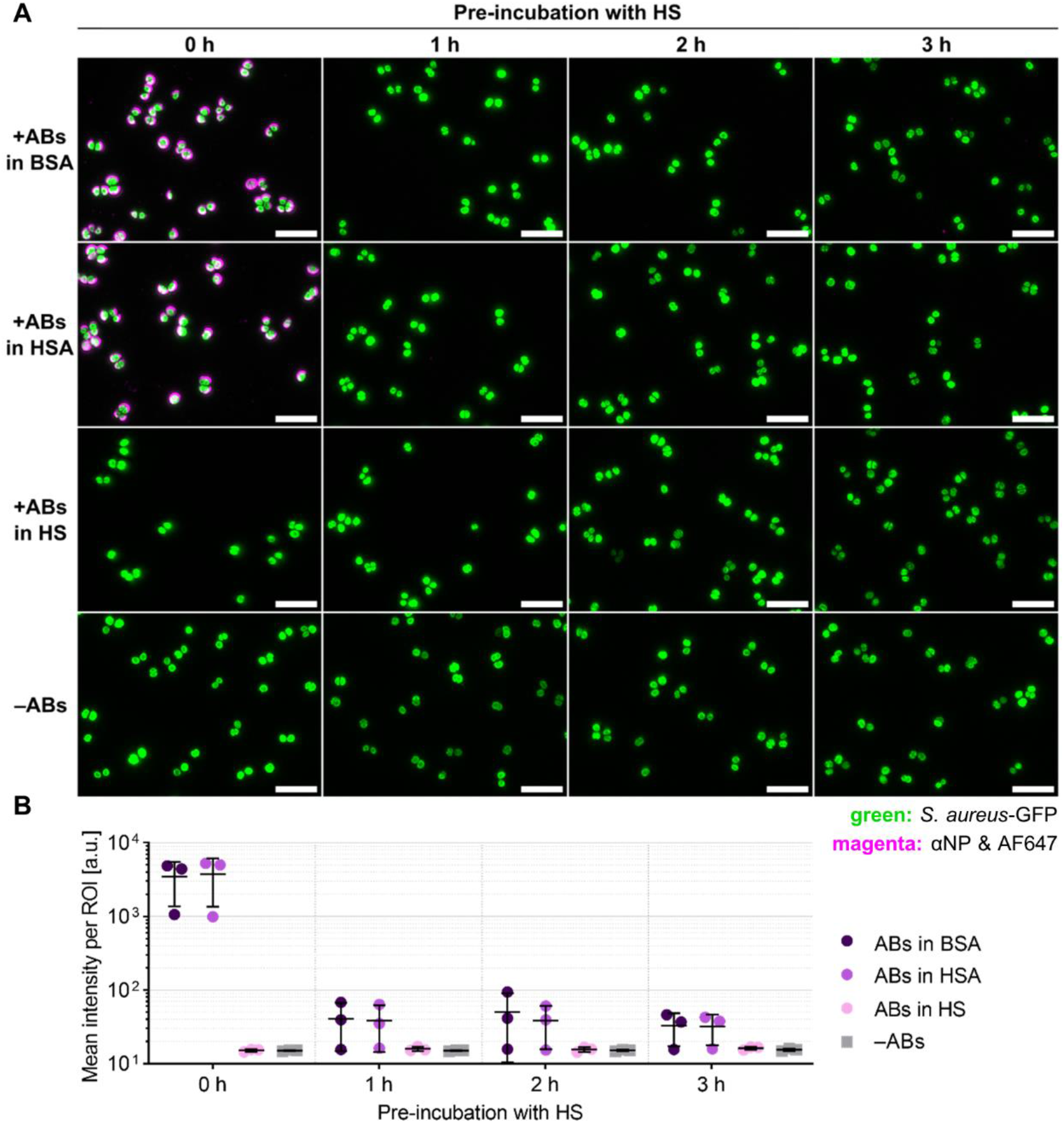
HS is a potent blocking agent and antibody diluent to prevent nonspecific antibody staining of *S. aureus*. Coverslip-coated GFP-expressing *S. aureus* was pre incubated with 100% HS for different incubation times (1-3 h) or not pre-incubated at all (0 h), and stained with the αNP/AF647 combination. Antibodies were diluted in 3% BSA, 3% HSA, or 50% HS. No-antibody controls (–ABs) were pre-incubated with 3% BSA. (A) Representative wide-field microscopy images. Scale bars = 5 µm. (B) Quantification of the nonspecific fluorescence as described in Supplemental Figure S2. Five images per condition were analyzed for each biological replicate. Depicted data points show medians of three biological replicates (Supplemental Figure S7); line and error bars show overall mean ± standard deviation. a.u.: arbitrary units; ABs: antibodies; BSA: bovine serum albumin; HS: human serum; HSA: human serum albumin; NP: nucleoprotein; ROI: region of interest.

### Validation of Blocking Strategies in *S. aureus*-infected Cells

Having evaluated the efficacy of αSpA and HS pre-incubation as well as antibody dilution in HS for blocking nonspecific antibody staining of *S. aureus*-coated on coverslips, we next sought to validate these approaches in a more biologically relevant context, in *S. aureus*-infected cells. Infected A549 cells were fixed, permeabilized, and subjected to the same blocking strategies, including pre incubation with αSpA (1:500 or 1:50 dilution in 3% BSA) or with 100% HS (Figure 5A-B), as well as varying HS incubation times (Figure 5C). Given its superior blocking efficacy for *S. aureus*-coated on coverslips (Figure 4B), antibodies in HS pre-incubated samples were diluted in 50% HS.

**Figure 5:**
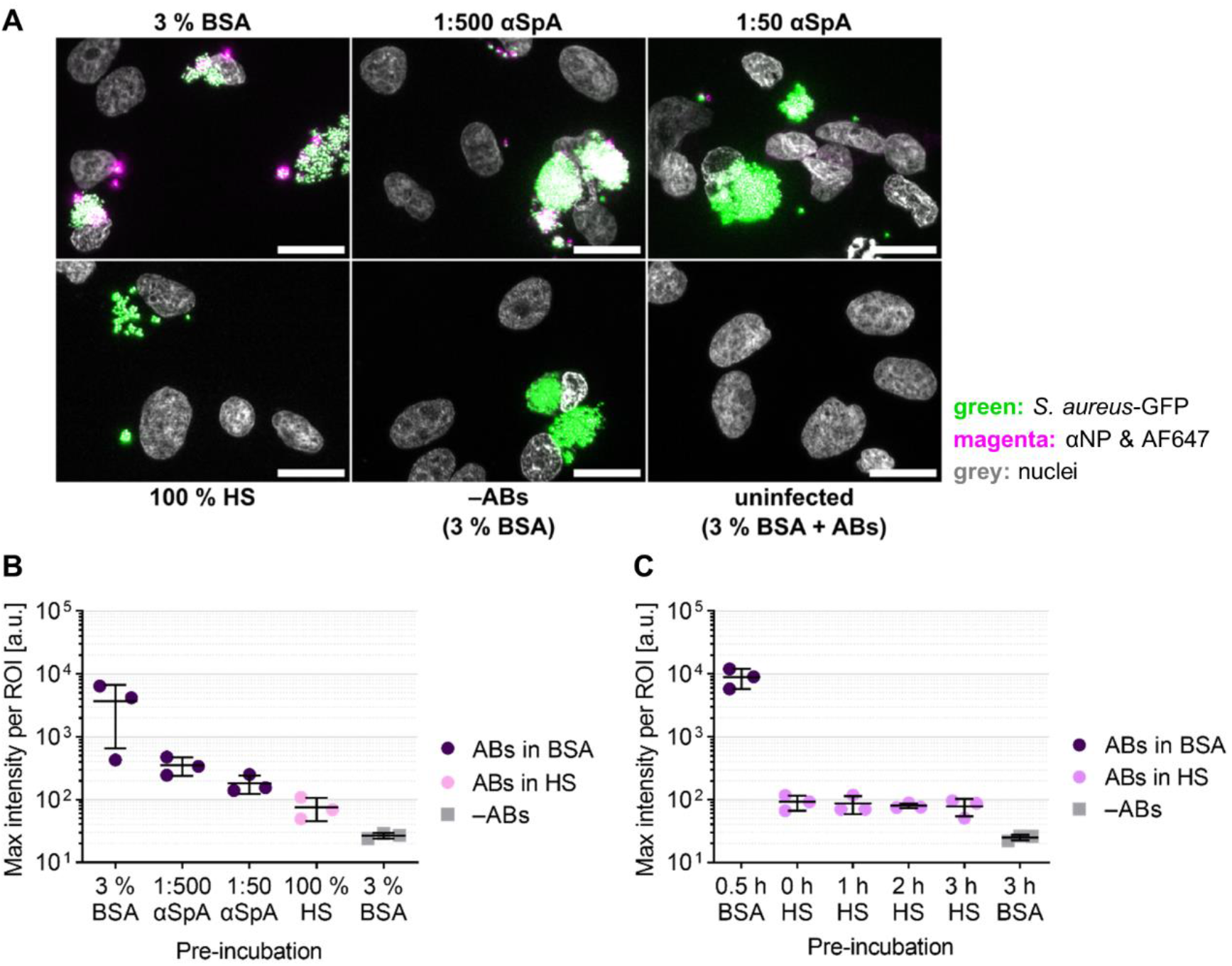
Validation of blocking strategies in *S. aureus*-infected cells. A549 cells were infected with GFP-expressing *S. aureus*. Prior to staining, cells were pre-incubated with BSA, αSpA, or HS. Cells were stained with the αNP/AF647 combination diluted in 3% BSA or 50% HS with additional Hoechst 33342 counterstaining. (A) Representative wide-field microscopy images. Of note, MIPs are shown in (A) for better contrast but segmentation and quantification were done in 3D. Scale bars = 20 µm. (B-C) Quantification of the nonspecific fluorescence as described in Supplemental Figure S4. Different agents for pre-incubation (B) and different incubation durations with HS (C) were compared. Five images per condition were analyzed for each biological replicate. Depicted data points show medians of three biological replicates (Figures S8-S9); line and error bars show overall mean ± standard deviation. a.u.: arbitrary units; ABs: antibodies; BSA: bovine serum albumin; HS: human serum; max: maximum; NP: nucleoprotein; ROI: region of interest; SpA: *staphylococcal* protein A.

Consistent with our previous findings, pre-incubation with either 1:50-diluted αSpA or with 100% HS effectively reduced nonspecific fluorescence, mirroring the trends observed on coverslip-coated *S. aureus*. Notably, pre-incubation with 100% HS appeared more effective than 1:50-diluted αSpA; however, since antibodies were diluted in 50% HS in the former condition but in 3% BSA in the latter, we cannot exclude the possibility that differences in antibody diluents contributed to the observed effect (Figure 5B). Furthermore, even in the absence of a dedicated pre-incubation step, antibody dilution in 50% HS once again resulted in a substantial reduction of nonspecific fluorescence, decreasing maximum intensities per ROI from the 10^4^ to the low 10^2^ a.u. range (Figure 5C).

While these results demonstrate that blocking strategies optimized in isolated *S. aureus* coated on coverslips translate well to infected cells, some residual fluorescence signal remained detectable in the intracellular environment. The lowest maximum intensities per ROI measured ranged from 50 to 120 a.u., whereas no-antibody controls showed maximum intensities of 20 to 30 a.u. per ROI. This discrepancy, which was not observed in coverslip-coated *S. aureus*, suggests that factors specific to the intracellular environment, such as bacterial clustering or interactions with the host cell, may contribute to low levels of residual nonspecific fluorescence. Therefore, careful selection of appropriate controls remains essential when performing immunofluorescence microscopy of *S. aureus*-infected cells.

Overall, with proper controls in place, pre-incubation with 100% HS together with antibody dilution in 50% HS provides an effective and broadly applicable method to mitigate SpA mediated nonspecific antibody labeling of *S. aureus* in immunofluorescence-based studies.

## Discussion

Immunofluorescence (IF) microscopy is a powerful tool to investigate host-pathogen interactions *in vitro*. However, the strong Fc-binding capacity of SpA presents a major obstacle for immunofluorescence studies in *S. aureus*-infected cells: SpA indiscriminately captures primary and secondary antibodies, irrespective of their antigen specificity. As a result, attempts to visualize, e.g., host cell factors are always confounded by nonspecific staining of the *S. aureus* surface, severely limiting quantitative analyses. In this study, we systematically evaluated two strategies to mitigate SpA-mediated interference. In both methods, the blocking agents, αSpA or HS, effectively saturated Fc-binding sites on SpA, thereby reducing nonspecific fluorescence. Notably, while αSpA pre-incubation lowered nonspecific signals by more than one order of magnitude, the most profound suppression was achieved by pre-incubation with and antibody dilution in HS. With the latter, blocking efficacy was nearly complete for *S. aureus*-coated on coverslips, where nonspecific signals were reduced to background levels. In *S. aureus*-infected cells, low levels of residual nonspecific fluorescence persisted, suggesting that additional optimization may further enhance the blocking efficacy in infected cells. Future refinements could include diluting αSpA in HS during the pre-incubation step or titrating HS as antibody diluent, as antibody dilution in concentrations higher than 50% HS may further improve suppression of SpA driven artifacts. Alternatively, pure Fcγ preparations could be tested for their inhibitory potential, as previous experiments revealed that Fcγ had a slightly stronger inhibitory effect on the binding of Fcγ to SpA than intact human IgG.^[20]^ Furthermore, a potential explanation for the observed difference in blocking efficacy between αSpA and HS lies in the distinct capacities of IgG molecules from different species to competitively bind to SpA.^[39, 54]^ Inganäs *et al*. demonstrated that IgG preparations from human, pig, dog, guinea pig, rhesus monkey, and rabbit were particularly effective inhibitors of SpA Fcγ binding, while IgG preparations from goat, sheep, mouse, rat, horse, and cow were markedly less potent.^[20]^ In line with these findings, Boyle *et al*. reported poor binding reactivity of most goat immunoglobulins to SpA.^[55]^ Therefore, the superior performance of HS in our experiments is likely attributable to its high content of human IgGs, in contrast to the αSpA that we applied, which was raised in goat.

Fc-mediated antibody binding by SpA is not only problematic for immunofluorescence microscopy but is a well-documented challenge across various immunoassays.^[44-46]^ Cronin *et al*. tested different blocking strategies as well as several factors that could affect the interaction between SpA and antibodies, including bacterial strain, heat killing, overnight storage, bacterial growth phase, and bacterial cell physiology, in the context of flow cytometry. They revealed that SpA-mediated antibody binding is highly strain-specific and that *S. aureus* with intact membranes and good cellular vitality exhibited a higher propensity for antibody binding. These findings underscore the importance of considering not only blocking strategies but also bacterial strain and physiological state. Moreover, it was demonstrated that a commercially available Fc receptor blocking reagent effectively mitigated SpA-mediated interference in flow cytometry, suggesting its potential applicability across different immunodetection methods.^[45]^

SpA consists of five homologous Ig binding domains^[56, 57]^ that do not only firmly bind to both the Fc and Fab regions of several antibody subtypes but that have also been shown to interact with host cell factors, such as tumor necrosis factor receptor 1,^[58]^ epithelial growth factor receptor,^[59]^ and von Willebrand factor (vWF).^[60, 61]^ These interactions might impose another level of interference in host-pathogen interaction studies in *S. aureus*-infected cells but similar blocking strategies using αSpA or IgG have successfully been employed to inhibit the SpA-vWF interaction^[60]^ indicating the versatility of these approaches.

Beyond blocking reagents, non-IgG based labeling poses an alternative strategy to circumvent SpA-mediated interference in immunofluorescence microscopy. One approach is to use antibody fragments devoid of Fc regions, such as monovalent antigen binding fragments with or without hinge region (Fab’/Fab), bivalent antigen binding fragments (F(ab’)2), single chain variable fragments (scFv), or custom-made nanobodies,^[62]^ in order to bypass the SpA-Fc interaction as stated on the webpages of some antibody suppliers.^[63, 64]^ Accordingly, Tan *et al*. reported no detectable antibody binding to fixed *S. aureus* on microscope slides when using secondary Fab’ fragments in a semi-quantitative direct immunofluorescence setup.^[47]^ However, others have shown that SpA also binds nonspecifically to the IgG Fab of mice,^[65]^ guinea pigs,^[66]^ and humans.^[67]^ Therefore, careful evaluation of undesired binding reactivities would be advisable for every antibody fragment. Alternatively, non-immunoglobulin-based probes, such as aptamers,^[68-70]^ offer highly promising solutions to circumvent nonspecific staining of SpA, but aptamers require specialized development and are nowadays still expensive, limiting their widespread applications in immunofluorescence microscopy.

Additionally, the use of *S. aureus* strains with inherently low SpA expression, such as E1369 or Wood 46,^[46, 71, 72]^ or culturing *S. aureus* on mannitol salt agar, which suppresses SpA production,^[40, 46, 73]^ represent possible strategies to avoid the nonspecific binding problem. Yet, nearly all clinically relevant isolates express SpA,^[74]^ making them difficult to bypass in infection models. Alternatively, genetically modified strains lacking SpA (Δ*spa*)^[60, 75]^ or strains expressing mutant SpA deficient in Fc-binding, like SpA_KKAA_^[76, 77]^ or SpA_KKE_/SpA_KKT_,^[78]^ could reduce nonspecific binding. However, genetic mutagenesis of *spa* raises concerns about bacterial pathogenicity and biological relevance, as it was shown that SpA-negative mutants were slightly less virulent than their SpA-positive counterparts,^[79]^ potentially limiting the applicability of such approaches for studying host-pathogen interactions. Additionally, SpA is not the only Fc-binding protein in *S. aureus*.^[80, 81]^ The *staphylococcal* binder of IgG (Sbi) contains two N-terminal IgG-binding domains that share some sequence similarity with the IgG-binding domains of SpA.^[81, 82]^ Therefore, a more general approach to block Fc-binding sites of both SpA and Sbi might yield better results than manipulating SpA itself.

## Conclusion

The confounding effects of SpA-mediated antibody binding in immunological detection methods have long been recognized within microbiology and immunology research communities, and strategies to mitigate this phenomenon have previously been described. However, this knowledge – and its practical implementation – has remained largely confined to a specific biological research context and is not always readily apparent to researchers approaching *S. aureus* from a biophysical or microscopy-driven perspective. As a result, SpA-mediated artifacts may go unrecognized or insufficiently controlled in imaging-based studies outside this traditional research domain.

Among the available countermeasures, HS emerges as a particularly effective and practical solution for immunofluorescence microscopy. Its ability to substantially reduce nonspecific fluorescence with minimal experimental modification, combined with its low cost and ease of implementation compared to engineered antibody fragments or genetically modified bacterial strains, makes HS a broadly accessible blocking strategy for imaging-focused laboratories. Nevertheless, irrespective of the blocking strategy chosen, blocking efficacy must always be carefully validated for each antibody combination and bacterial strain before experimental application. Factors such as antibody affinity, bacterial strain variability, and experimental conditions can all influence SpA-mediated interference, necessitating thorough pre-evaluation to assure reliable results. For optimal performance, pre-testing with *S. aureus* coated on coverslips provides a practical approach to assess and fine tune blocking conditions before applying the method to more complex infection models.

Overall, our findings establish HS as a simple yet highly effective reagent for preventing SpA-driven artifacts in immunofluorescence microscopy. We provide experimentally validated and quantitative guidance intended to facilitate the broader adoption of robust blocking strategies within the microscopy and biophysics communities.

## Materials and Methods

Detailed experimental descriptions can be found in the Supporting Information.

## Supporting Information

The authors have cited additional references within the Supporting Information.^[83-87]^

## Acknowledgements

We thank Dr. Bianca Hoffman (Leibniz Institute for Natural Product Research and Infection Biology, Jena, Germany) for her great support with writing macros for quantifying the nonspecific labeling of *S. aureus*. We thank Dr. Daniel Lopez (Spanish National Research Council CNB-CSIC, Madrid, Spain) for fruitful discussions on general bacteria labeling. We thank the Microverse Imaging Center (Dr. Aurélie Jost and Dr. Patrick Then) for providing microscope facility support for data acquisition. The authors greatly acknowledge financial support by the Deutsche Forschungsgemeinschaft (DFG, German Research Foundation; Germany’s Excellence Strategy – EXC 2051 – Project-ID 390713860; SFB 1278/2 PolyTarget-316213987-A04, D01, D02, Z01; GRK M-M-M: GRK 2723/1 – 2023 – ID 44711651; GRK PhInt: GRK 3014/1; Instrument funding modular STED INST 1757/25-1 FUGG; instrument funding ID 460889961 multi-photon laser scanning device), the State of Thuringia (TMWWDG), the Leibniz Association (Leibniz ScienceCampus InfectoOptics Jena financed by the funding line Strategic Networking of the Leibniz Association, project number W8/2018; and Leibniz Collaborative Excellence Programme, project AMPel – project number K548/2023), the Free State of Thuringia (TAB; AdvancedSTED / FGZ: 2018 FGI 0022; Advanced Flu-Spec / 2020 FGZ: FGI 0031; Multi-XUV / 2023 FGR 0054), the innovation program by the German BMWi (ZIM; project 16KN070934 / Lab-on-a-chip FCS-Easy, project 16KN070967 / Lab-on-a-chip SMARTIES). Further, this work is supported by the BMFTR (Federal Ministry of Research, Technology and Space), funding program LIVE2QMIC (FGZ: 13N15956) as well as Photonics Research Germany (FKZ: 13N15713 / 13N15717) and is integrated into the Leibniz Center for Photonics in Infection Research (LPI). The LPI initiated by Leibniz-IPHT, Leibniz-HKI, UKJ and FSU Jena is part of the BMFTR national roadmap for research infrastructures. A part of the project on which these results are based was funded by the Free State of Thuringia under the number 2018 IZN 0002 (Thimedop) and co-financed by funds from the European Union within the framework of the European Regional Development Fund (EFRE).

