## Supporting Information for "Overcoming Protein A-driven Nonspecific Antibody Staining of *S. aureus* in Immunofluorescence Microscopy"

#### This PDF file includes:

Experimental models
Methods
Materials
Supplementary figures
References

### Experimental models

#### Cell lines

A549 cells were cultured in DMEM with high glucose and L-glutamine but without sodium pyruvate supplemented with 10% fetal constance (FC) at 37 °C, 5% $CO_2$, 95% relative humidity (rH) and cells were passaged every 4 to 5 days.

#### Bacteria

Glycerol stocks (25% glycerol) of GFP-expressing *S. aureus* USA300 were stored at -80 °C. Overnight cultures (ONCs) were directly prepared from glycerol stocks as follows: *S. aureus* was recovered from glycerol stocks using a pipette tip and cultivated in 50 mL Erlenmeyer flasks containing 5 mL BHI medium supplemented with 30 µg $mL^{-1}$ chloramphenicol. Bacteria were grown at 37 °C and 160 rpm overnight. ONCs were transferred into 15 mL falcons, centrifuged (3000 xg, 5 min, 4 °C), and washed once with PBS. After another centrifugation (as before), supernatant was removed and bacteria were resuspended in 5 mL PBS. $OD_{600}$ was measured, adjusted to ≈ 0.7 by diluting the ONC with PBS, and the final $OD_{600}$ was determined.

### Methods

#### Buffers

Phosphate-buffered saline (PBS): 137 mM NaCl, 2.7 mM KCl, 8 mM $Na_2HPO_4$, 1.5 mM $KH_2PO_4$ in dd$H_2O$, pH 7.4.
Invasion medium (DMEM/INV): 1% HSA, 1 mM HEPES in DMEM.
Infection medium (DMEM/INF): 1 mM $MgCl_2$, 0.9 mM $CaCl_2$, 0.21% (w/v) BSA in DMEM.

#### *S. aureus* coating on coverslips

For STED, #1.5H coverslips were used while #1 coverslips were utilized for wide-field microscopy. In both cases, coverslips were coated with 0.01% (w/v) poly-L-lysine in 24-well plates for 5 min and washed once with PBS. After PBS removal, coverslips were dried at room temperature (RT) and 300 µL *S. aureus* ONC with $OD_{600}$ ≈ 0.7 were added for 10 min. After another PBS wash, bacteria were fixed in 3.7% formaldehyde for 15 min at RT, washed thrice with PBS, and stored in PBS at 4 °C until staining.

#### *S. aureus* infection of A549 cells

1.5 * $10^5$ A549 cells per well were seeded on #1 coverslips in 500 µL DMEM in 24-well plates. 24 h after seeding, cells were washed once with PBS and infected with GFP-expressing *S. aureus* USA300 at a multiplicity of infection of 50 (MOI 50) whereat the ONC was diluted in DMEM/INV to a total volume of 250 µL per well. Two assumptions were made to calculate the *S. aureus* ONC volume ($V_{ONC}$): (1) The A549 cell number per well roughly doubled during the 24 h since seeding. (2) At $OD_{600} = 1$, *S. aureus* has a titer of 5 * $10^5$ cfu $\mu L^{-1}$. With this, $V_{ONC}$ was calculated as follows:

$$V_{ONC} = \frac{1}{OD_{600}} * \frac{MOI * cell\ number}{titer\ at\ OD_{600} = 1} = \frac{1}{\mathrm{OD}_{600}} * \frac{50\ \mathrm{cfu} * \frac{3 * 10^5}{\mathrm{well}}}{5 * 10^5\ \frac{\mathrm{cfu}}{\mu\mathrm{L}}} = \frac{30}{\mathrm{OD}_{600}} \frac{\mu\mathrm{L}}{\mathrm{well}}$$

Inocula were prepared in DMEM/INV accordingly and 250 µL per well were added. Cells were shortly centrifuged (200 xg, 2 min, RT), incubated at 37 °C, 5% $CO_2$, 95% rH for 3 h, and washed once with PBS. To remove the remaining extracellular bacteria, cells were incubated with 2 µg $mL^{-1}$ lysostaphin diluted in DMEM/10% FC for 20 min as above. Cells were washed once with PBS and kept in DMEM/INF at 37 °C, 5% $CO_2$, 95% rH until fixation at 9.5 h post-infection. Cells were fixed in 3.7% formaldehyde for 15 min at RT, washed thrice with PBS, and stored in PBS at 4 °C until staining.

##### Antibody staining for STED microscopy

All following steps were performed at RT. Antibodies were always diluted in 3% BSA in PBS (BSA/PBS). Fixed coverslip-coated *S. aureus* was blocked with 3% BSA/PBS for 30 min and washed once with PBS. The bacteria were then incubated with goat-α-SpA (1:50) or mouse-α-NP (1:300) for 1 h. After three PBS wash steps, bacteria were stained with donkey-α-goat-AF568 (1:300), chicken-α-goat-AF647 (1:300), donkey-α-mouse-AF568 (1:300), or donkey-α-mouse-AF647 (1:300) for 30 min in darkness. After three PBS wash steps, coverslips were mounted on objective slides using fluorescence mounting medium and stored at 4 °C until imaging.

##### Pre-incubation with αSpA and antibody staining

All following steps were performed at RT. *S. aureus*-infected cells were permeabilized with 0.1% Triton X-100 for 20 min and cells were washed thrice with PBS. For *S. aureus*-coated on coverslips, permeabilization was not required. Both sample types (*S. aureus* coated on coverslips and *S. aureus*-infected cells) were blocked with 3% BSA/PBS for 30 min and washed once with PBS. In the following, all antibodies were prepared in 3% BSA/PBS. The samples were pre-incubated with increasing amounts (1:1000 to 1:50) of goat-α-SpA or kept in 3% BSA/PBS (no-αSpA and no-antibody controls) for 1 h. Samples were washed thrice with PBS and incubated with mouse-α-NP (1:300) or kept in 3% BSA/PBS (no-antibody control) for 1 h. After three PBS wash steps, the samples were stained for 30 min in darkness with donkey-α-mouse-AF647 (1:500) and, in the case of infected cells, together with Hoechst 33342 (1:2000). No-antibody controls were only stained with Hoechst 33342 (1:2000; infected cells) or kept in 3% BSA/PBS (coverslip-coated *S. aureus*) for 30 min in darkness. After three PBS wash steps, samples were mounted and stored as above.

For coverslip-coated *S. aureus*, the blocking efficacy of αSpA was tested for various antibody combinations (Figure 3C). Bacteria were pre-incubated with αSpA (1:500 or 1:50) and stained as described above. Table S1 indicates all tested antibodies and their dilutions.

**Table S1:** Antibody combinations and their respective dilutions underlying data in Figure 3C.

| Combination | Primary antibody | Secondary antibody |
|---|---|---|
| E-Cad/Cy5 | rabbit-α-E-Cad (1:150) | goat-α-rabbit-Cy5 (1:150) |
| ERK2/Cy5 | mouse-α-ERK2 (1:50) | goat-α-mouse-Cy5 (1:150) |
| M1/Cy5 | mouse-α-M1 (1:100) | goat-α-mouse-Cy5 (1:150) |
| PB1/Cy5 | rabbit-α-PB1 (1:100) | goat-α-rabbit-Cy5 (1:150) |
| NS1/AF647 | mouse-α-NS1 (1:50) | donkey-α-mouse-AF647 (1:300) |
| NP/Cy3 | mouse-α-NP (1:300) | goat-α-mouse-Cy3 (1:300) |
| p38α/Cy3 | rabbit-α-p38α (1:50) | goat-α-rabbit-Cy3 (1:300) |
| p50/AF568 | rabbit-α-p50 (1:100) | donkey-α-rabbit-AF568 (1:300) |

α: anti; E-Cad: E-Cadherin; ERK2: extracellular signal-regulated kinase 2; M1: matrix protein 1; NP: nucleoprotein; NS1: non-structural protein 1; p38α: Mitogen-activated protein kinase 14; p50: p50 subunit of NFκB; PB1: polymerase basic protein 1.

##### Pre-incubation with HS and antibody staining

All incubation steps were performed at RT. *S. aureus* coated on coverslips was pre-incubated with 100% HS for 1-3 h or not pre-incubated (0 h) and washed once with PBS. Bacteria were then treated with mouse-α-NP (1:300) diluted in 50% HS/PBS, 3% BSA/PBS, or 3% HSA/PBS for 1 h, while no-antibody controls were kept in3 % BSA/PBS. After three PBS wash steps, bacteria were stained in darkness using donkey-α-mouse-AF647 (1:500) diluted as before for 30 min. No-antibody controls were again kept in 3% BSA/PBS. After three PBS wash steps, samples were mounted and stored as above.

For *S. aureus*-infected cells, the protocol was slightly different. Cells were permeabilized with 0.1% Triton X-100 for 20 min and washed thrice with PBS. Cells were pre-incubated with 100% HS for 1-3 h (HS pre-incubation), with 3% BSA/PBS for 0.5 h (positive control), with 3% BSA/PBS for 3 h (no-antibody control), or not pre-incubated (0 h). Cells were washed once in PBS and incubated for 1 h with mouse-α-NP (1:300) diluted in either 50% HS/PBS (HS pre-incubation) or in 3% BSA/PBS (positive control), while the no-antibody control was kept in 3% BSA/PBS. After three PBS wash steps, cells were stained for 30 min in darkness with donkey-α-mouse-AF647 (1:500) diluted as before and with

Hoechst 33342 (1:2000), while the no-antibody control was only stained with Hoechst 33342 (1:2000). After three PBS wash steps, samples were mounted and stored as above.

**Wide-field fluorescence microscopy**

Z-stack images were acquired with an Axio Observer.Z1 microscope (Zeiss, 431007-9901-000) equipped with ApoTome.2 (Zeiss, 423667-8224-000), Axiocam 503 mono camera (Zeiss, 26559 0000 000), HXP 120 V light source (Leistungselektronik Jena GmbH, LQ-HXP120-CAN-z-v), and Zen blue. Coverslip-coated *S. aureus* was imaged using a 63x/NA 1.40 (Zeiss, 420782-9900-799) or a 100x/NA 1.30 (Zeiss, 420490-9900-000) oil immersion objective with Immersol™ 518 F. *S. aureus*-infected cells were imaged using a 20x/NA 0.8 (Zeiss, 420650-9902-000) air objective. For image acquisition, the following filter sets were applied: 50 (Zeiss, 488050 9901 000) for Cy5 and AF647, 43 (Zeiss, 000000-1114-101) for Cy3 and AF568, 38 HE (Zeiss, 489038-9901-000) for GFP, and 49 (Zeiss, 488049-9901-000) for Hoechst 33342. Z-stacks were recorded as 14-bit images and, after acquisition, images were directly deconvolved in Zen blue with phase errors correction and deconvolution strength 5. In case of *S. aureus*-infected cells, deconvolved z-stacks were exported for further processing. For coverslip-coated *S. aureus*, images were maximum intensity projected in Zen blue, prior to the export. For image display, the dynamic ranges of 2- or 3-channel MIPs were adjusted using Fiji[1] as described in Table S2. Of note, in all figures, MIP insets (as depicted in Figure 2) are displayed for better recognizability, but quantification is always based on the entire images.

**Table S2:** Displayed dynamic ranges of wide-field images in Figures 2-5 and Figure S1. For image display, MIPs were loaded into Fiji[1] as 16-bit images and the displayed dynamic ranges for the individual channels were set as follows.

| Figure | Dye | Saturation set to | Displayed dynamic range |
|---|---|---|---|
| 2A | AF647 | - | 0 – 16000 |
| | GFP | - | 0 – 16000 |
| 2C | AF647 | - | 10 – 1400 |
| | GFP | - | 50 – 1000 |
| | Hoechst 33342 | 0.2 | - |
| 3A | AF647 | - | 50 – 10000 |
| | GFP | - | 50 – 12000 |
| 4A | AF647 | - | 0 – 8000 |
| | GFP | - | 0 – 8000 |
| 5A | AF647 | - | 30 – 1600 |
| | GFP | - | 30 – 1200 |
| | Hoechst 33342 | 0.2 | - |
| S1A | Cy5/AF647 | 0.35[a] | 12 – 724[b] |
| | GFP | 0.35 | - |
| S1B | Cy3/AF568 | 0.35[a] | 19 – 1365[b] |
| | GFP | 0.35 | - |
| S1C | Cy5/AF647 | 0.35[a] | 9 – 63[b] |
| | GFP | 0.35 | - |
| S1D | Cy3/AF568 | 0.35[a] | 16 – 808[b] |
| | GFP | 0.35 | - |

[a]Saturation of the red or orange channels was set to 0.35 for all but the no-antibody control images in this figure.
[b]For the no-antibody control images, the displayed dynamic range was aligned with the smallest dynamic range of all other images in this figure.

#### STED microscopy

STED microscopy was performed using an Abberior STEDYCON (Abberior Instruments, revision 2) on an Olympus inverted microscope body (Olympus, IX83) equipped with STEDYCON smart control. Images were acquired using a 100x/NA 1.45 oil immersion objective (Olympus, UPLXAPO100XO) with type F immersion oil. Images were recorded by exciting AF568 with 561 nm, AF647 with 640 nm, and GFP with 488 nm laser lines with powers as indicated in Table S3. All excitation lasers were pulsed diode lasers with a repetition rate of 40 MHz and a pulse duration of <150 ps. A STED beam at 775 nm has been used to deplete the orange or red laser line. The fixed pinhole diameter in the intermediate image plane was 66.66 µm. Multichannel images were recorded in line-interleaved scanning mode with up to four line accumulations (Table S3), a pixel dwell time of 10 µs, and a fixed pixel size of 40 nm. Photons were collected by single-photon counting avalanche photodiodes (Excelitas Technologies, SPCM-AQRH-13-FC) equipped with appropriate filters (500-550 nm, 580-630 nm, and 650-700 nm). For orange and red channels, photons were collected between 1 to 7 ns after the excitation pulse. All STED images shown are raw data.

**Table S3:** Acquisition settings for STED images shown in Figure 1.

| Position in Figure 1 (AB combination) | Excitation laser | | STED laser | | Line acc. | Detector | |
|---|---|---|---|---|---|---|---|
| | *λ* [nm] | *P* [µW] | *λ* [nm] | *P* [mW] | | *λ* [nm] | gate [ns] |
| Top left (SpA/AF568) | 561 | 0.4 | 775 | 124.8 | 3 | 580-630 | 1-7 |
| | 488 | 0.1 | - | - | | 500-550 | - |
| Bottom left (SpA/AF647) | 640 | 1.0 | 775 | 124.9 | 1 | 650-700 | 1-7 |
| | 488 | 0.2 | - | - | | 500-550 | - |
| Top right (NP/AF568) | 561 | 0.3 | 775 | 121.1 | 4 | 580-630 | 1-7 |
| | 488 | 0.1 | - | - | | 500-550 | - |
| Bottom right (NP/AF647) | 640 | 1.6 | 775 | 60.4 | 1 | 650-700 | 1-7 |
| | 488 | 0.3 | - | - | | 500-550 | - |

AB: antibody; acc.: accumulations; AF: Alexa Fluor; NP: nucleoprotein; SpA: *staphylococcal* protein A; STED: stimulated emission depletion.

#### Image processing and quantification of nonspecific antibody labeling of coverslip-coated *S. aureus*

2-channel MIPs (green channel: GFP-expressing *S. aureus*; red or orange channel: nonspecific fluorescence) were loaded into Fiji[1] as 16-bit images and processed as follows (Figure S2): Channels were split and processed separately. The green MIP (GFP-expressing *S. aureus*) was tresholded using Renyi's entropy. The resulting binary image was processed with fill holes, close-, dilate, and adjustable watershed with a tolerance of 0.08 for 63x images and a tolerance of 0.1 for 100x images. From the binary image, ROIs were extracted using Fiji's inbuilt particle analyzer with a size setting of 0.5-infinity for both 63x and 100x images but no restrictions for circularity. Mean red or orange intensities (nonspecific signals derived from the antibody staining) within each ROI were measured with three decimal places. In total, 5 images per condition were analyzed, unless stated differently. Mean red or orange intensities per ROI were plotted for individual experiments and the median ± interquartile range of the mean red or orange intensities per ROI were determined for each condition. Once data from several biological replicates were obtained, these median values were plotted and mean ± standard deviation were calculated.

#### Image processing and quantification of nonspecific antibody labeling of *S. aureus* in infected A549 cells

3-channel z-stacks (green channel: GFP-expressing *S. aureus*; red channel: nonspecific fluorescence; blue channel: nuclei) were loaded into Fiji[1] as 16-bit images and processed as follows (Figure S4): Channels were split and the red and green channels were saved separately. The blue channel was disregarded during quantification since nuclei were only stained for visualization. The green channel

(GFP-expressing *S. aureus*) was maximum intensity projected and the resulting MIP was thresholded using Renyi's entropy. From this thresholding, the lower threshold value was extracted and saved for later 3D segmentation. Next, the original green z-stack was processed with a 3D maximum filter with a radius of 2 and MorphoLibJ's[2] 3D ball-shaped dilation with a radius of 3. The processed green z-stack was then thresholded using the 3D ROI manager[3] entering the beforehand extracted lower threshold value as lower and 65535 (maximum value for 16-bit images) as upper border. Mean and maximum red intensity (nonspecific signals derived from the antibody staining) within each ROI of an image were measured. In total, 5 images per condition were analyzed, unless stated differently. Maximum red intensities per ROI were plotted for individual experiments, unless stated otherwise, and the median ± interquartile range were determined for each condition. Once data from several biological replicates were obtained, these median values were plotted and mean ± standard deviation were calculated.

#### Materials

All reagents, experimental models, software, and consumables used for this study are listed in Table S4.

**Table S4:** Reagents and resources used in this study.

| REAGENT OR RESOURCE | SOURCE | IDENTIFIER |
|---|---|---|
| **Antibodies** | | |
| Chicken-α-goat-AF647 | Invitrogen | A-21469 |
| Donkey-α-goat-AF568 | Invitrogen | A-11057 |
| Donkey-α-mouse-AF568 | Invitrogen | A-10037 |
| Donkey-α-mouse-AF647 | Invitrogen | A-31571 |
| Donkey-α-rabbit-AF568 | Invitrogen | A-10042 |
| Goat-α-mouse-Cy3 | Dianova | 115-165-003 |
| Goat-α-mouse-Cy5 | Dianova | 115-175-146 |
| Goat-α-rabbit-Cy3 | Dianova | 111-165-003 |
| Goat-α-rabbit-Cy5 | Dianova | 111-175-144 |
| Goat-α-SpA | Rockland | 100-1177 |
| Mouse-α-ERK2 | Santa Cruz | sc-1647 |
| Mouse-α-M1 | BioRad | MCA401 |
| Mouse-α-NP | Bio-Rad | MCA400 |
| Mouse-α-NS1 | Institute of Molecular Virology Münster, University of Münster, Germany | clone NS1-23-1[4] |
| Rabbit-α-E-Cad | Cell Signaling Technology | 3195 |
| Rabbit-α-p38α | Santa Cruz | sc-535 |
| Rabbit-α-p50 | Santa Cruz | sc-8414 |
| Rabbit-α-PB1 | GeneTex | 125923 |
| **Bacterial strains** | | |
| GFP-expressing *S. aureus* USA300 | Jordan *et al*.[5] | N/A |
| **Chemicals, peptides, and recombinant proteins** | | |
| Bovine serum albumin (BSA) | Carl Roth | 9401.3 |
| Brain-Heart-Infusion (BHI) medium | Carl Roth | X916.1 |
| $CaCl_2$ | Carl Roth | CN93.1 |
| Chloramphenicol | Carl Roth | 3886.2 |
| Dulbecco's Modified Eagle Medium (DMEM) | Anprotec | AC-LM-0012 |
| Fetal Constance (FC) | Anprotec | AC-SM-0190 |
| Formaldehyde | Carl Roth | 7398.1 |

**Table S4 (continued):** Reagents and resources used in this study.

| REAGENT OR RESOURCE | SOURCE | IDENTIFIER |
|---|---|---|
| **Chemicals, peptides, and recombinant proteins (continued)** | | |
| Glycerol | Carl Roth | 3783.3 |
| HEPES | Lonza | BE17-737E |
| Hoechst 33342 | Sigma-Aldrich | 14533-100MG |
| Human serum (HS) | Merck | S1 |
| Human serum albumin (HSA) | Octapharma | 5400949 |
| KCl | Carl Roth | 6781.9 |
| $KH_2PO_4$ | Carl Roth | 3904.2 |
| Lysostaphin | WAK-Chemie Medical | WAK-LSPN-50 |
| $MgCl_2$ | Sigma-Aldrich | M2393 |
| NaCl | ITW Reagents | A2942,1000 |
| $Na_2HPO_4$ | ITW Reagents | 131679.1210 |
| Poly-L-lysine | Merck | A-005-C |
| Triton X-100 | Sigma-Aldrich | X100 |
| **Experimental models: Cell lines** | | |
| A549 cells | American Type Culture Collection (ATCC) | CCL-185 |
| **Software and algorithms** | | |
| Excel | Microsoft | Version 2016 |
| Fiji Is Just ImageJ (Fiji)[1] | National Institutes of Health | Version 2.16.0/1.54p |
| GraphPad Prism | GraphPad Software Inc | Version 8.4.3 |
| STEDYCON smart control | Abberior Instruments | Firmware 7.1.53 |
| Zen blue | Zeiss | Version 2.6 |
| **Other** | | |
| #1 coverslips (130-160 µm thickness) | Carl Roth | P231.1 |
| #1.5H coverslips (165-175 µm thickness) | Marienfeld | 0117520 |
| Fluorescence mounting medium | Dako | S3023 |
| ImmersolTM 518 F | Zeiss | 444960-0000-000 |
| Objective slides | Epredia | AA00000112E01MNZ10 |
| Type F immersion oil | Olympus | IMMOIL-F30CC |

α: anti; AF: Alexa Fluor; Cy3/5: Cyanine 3/5; E-Cad: E-Cadherin; ERK2: extracellular signal-regulated kinase 2; GFP: green fluorescent protein; M1: matrix protein 1; NP: nucleoprotein; NS1: non-structural protein 1; p38α: Mitogen-activated protein kinase 14; p50: p50 subunit of NFκB; PB1: polymerase basic protein 1; SpA: *staphylococcal* protein A.

#### Supplementary figures

##### Figure S1

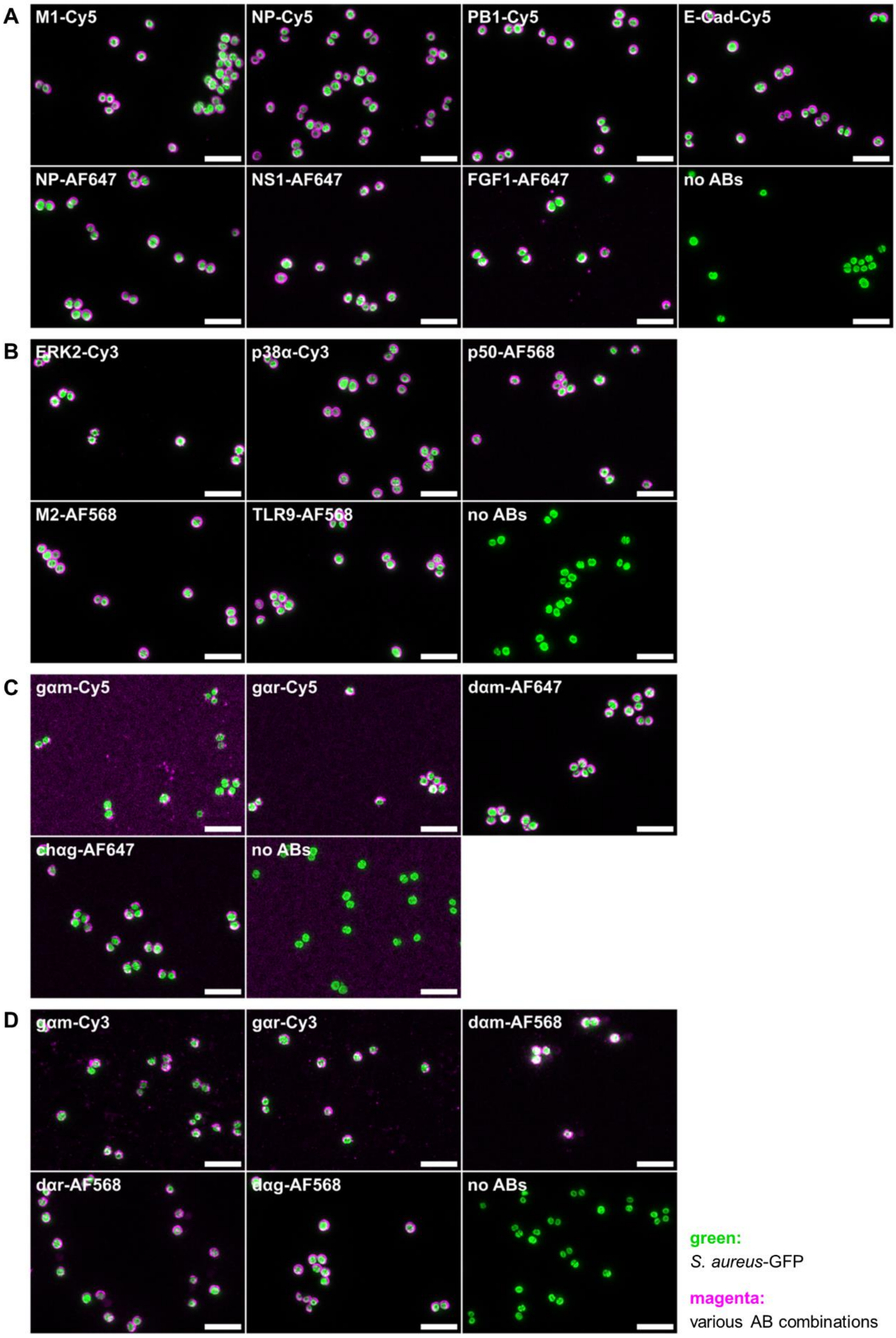


**Figure S1:** *S. aureus* is nonspecifically stained by various antibody combinations. (A-D) Wide-field images of GFP-expressing *S. aureus*. Coverslip-coated *S. aureus* was stained with various antibody combinations (A-B), only with secondary antibodies (C-D), or without antibodies (no ABs, A-D). Primary antibody targets included mammalian and viral antigens. Secondary antibodies were conjugated to different red (A, C) or orange (B, D) chromophores. Scale bars = 5 µm. AB: antibody; chαg: chicken-anti-goat; dαm: donkey-anti-mouse; dαr: donkey-anti-rabbit; dαg: donkey-anti-goat; E-Cad: E-Cadherin; ERK2: extracellular signal-regulated kinase 2; gαm: goat-anti-mouse; gαr: goat-anti-rabbit; M1: matrix protein 1; NP: nucleoprotein; NS1: non-structural protein 1; p38α: mitogen-activated protein kinase 14; p50: p50 subunit of NFκB; PB1: polymerase basic protein 1.

#### Figure S2

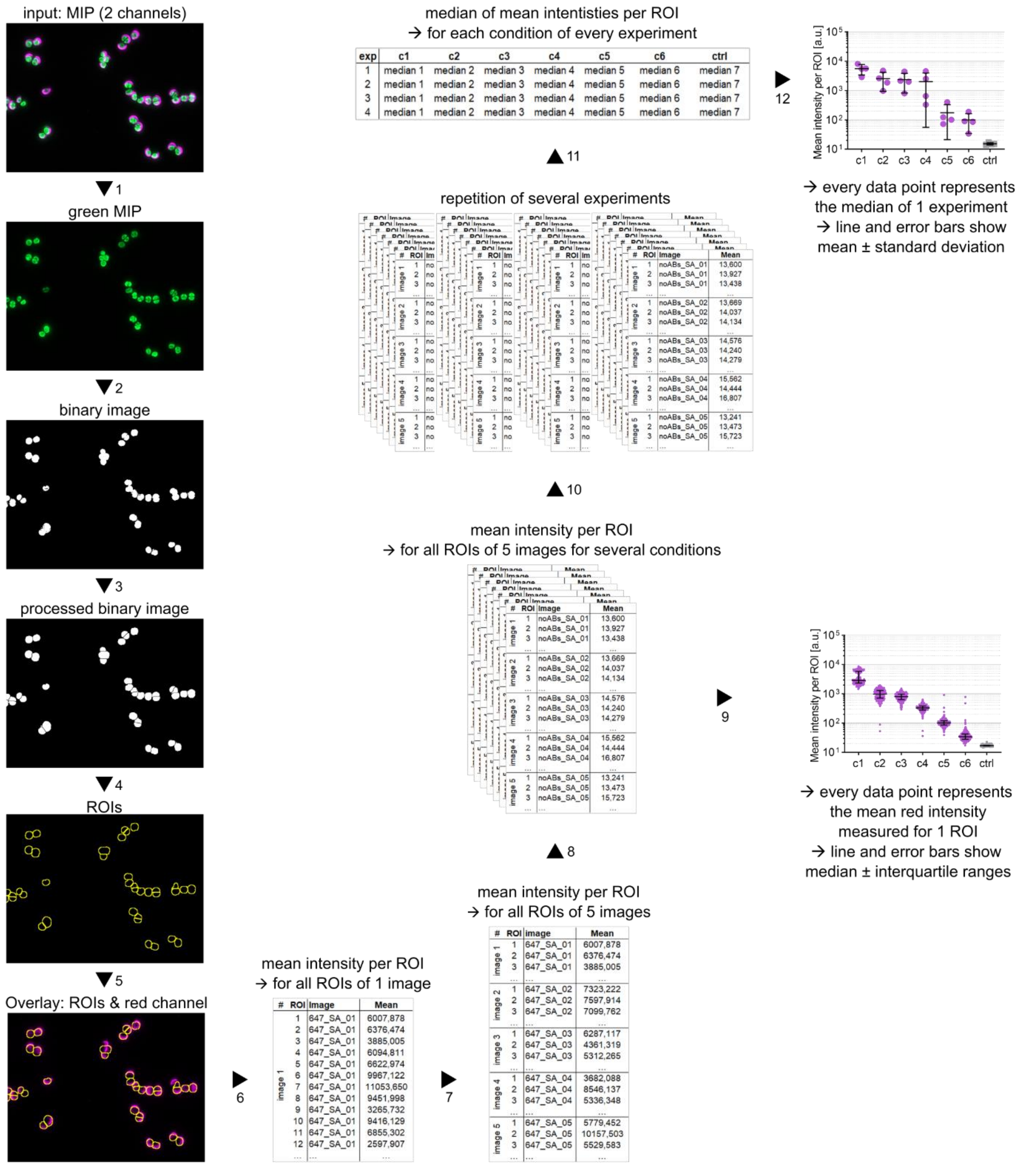


**Figure S2:** Image analysis pipeline for the quantification of nonspecific antibody staining of *S. aureus* coated on coverslips. 2-channel z-stacks were deconvolved and maximum intensity projected using Zen blue. Resulting MIPs were further processed and quantified in Fiji[1] as follows. 1: Extraction of the green channel. 2: Thresholding of the green channel using Renyi's entropy. 3: Processing of the binary image with fill holes, close-, dilate, and adjustable watershed. 4: Extraction of ROIs from the binary image. 5: Overlay of ROIs with the second channel (red or orange). 6: Measurement of the mean red intensity within each ROI. 7: Repetition of steps 1 to 6 for, in total, five images per condition. 8: Repetition of steps 1 to 7 for all conditions of an experiment. 9: To analyze individual experiments: Plotting of mean red intensities per ROI of five images for each condition. 10: For biological replicates: Repetition of steps 1 to 9 for each replicate. 11: To combine all biological replicates: Determination of the median of all mean red intensities per ROI for each condition and biological replicate. 12: Plotting of these median values, which results in one data point per condition and biological replicate. a.u.: arbitrary units; c: condition; ctrl: control; exp: experiment; MIP: maximum intensity projection; ROIs: regions of interest.

**Figure S3**

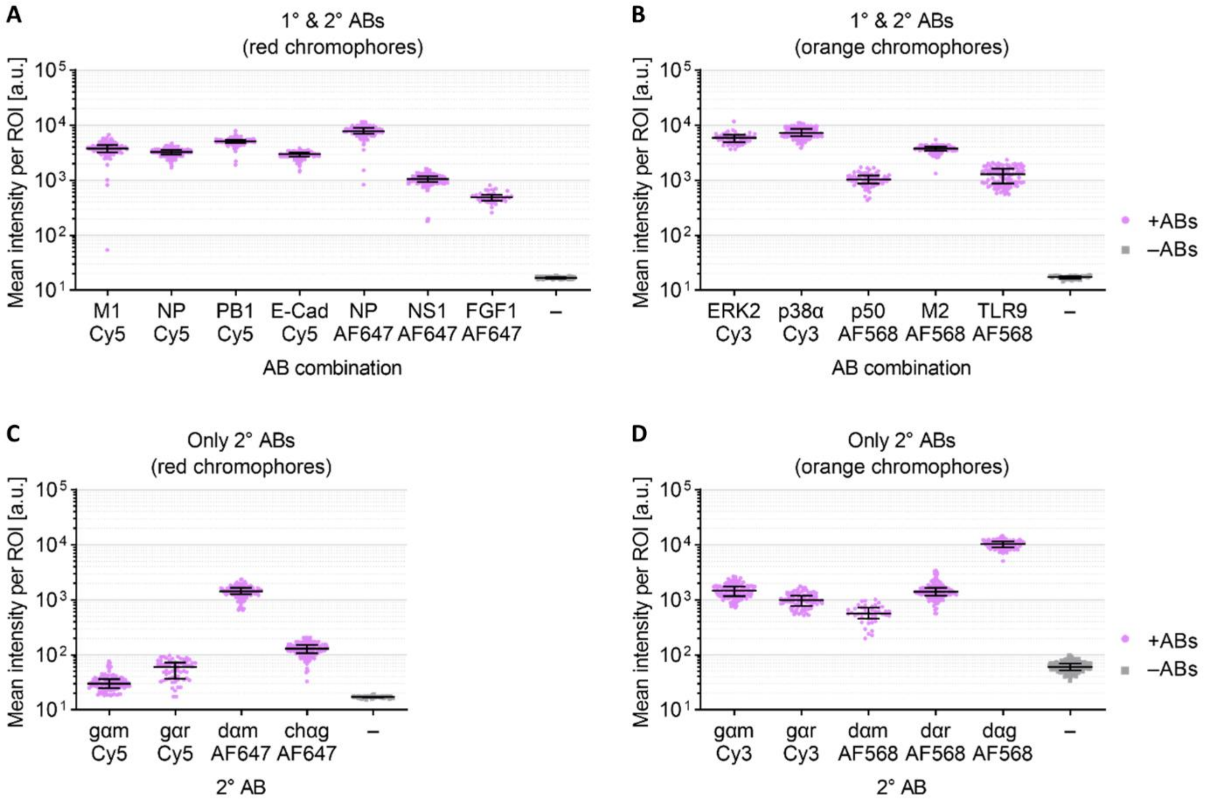


**Figure S3:** Quantification of nonspecific staining of *S. aureus* coated on coverslips for various antibody combinations. (A-D) Quantification corresponding to the experiments shown in Figure S1. Quantification was done as described in Figure S2, steps 1-9. Two images per condition were analyzed. Line and error bars depict median ± interquartile range. 1°: primary; 2°: secondary; a.u.: arbitrary units; ABs: antibodies; chαg: chicken-anti-goat; dαm: donkey-anti-mouse; dαr: donkey-anti-rabbit; dαg: donkey-anti-goat; E-Cad: E-Cadherin; ERK2: extracellular signal-regulated kinase 2; FGF1: fibroblast growth factor 1; gαm: goat-anti-mouse; gαr: goat-anti-rabbit; M1: matrix protein 1; M2: matrix protein 2; NP: nucleoprotein; NS1: non-structural protein 1; p38α: mitogen-activated protein kinase 14; p50: p50 subunit of NFκB; PB1: polymerase basic protein 1; TLR9: Toll-like receptor 9.

#### Figure S4

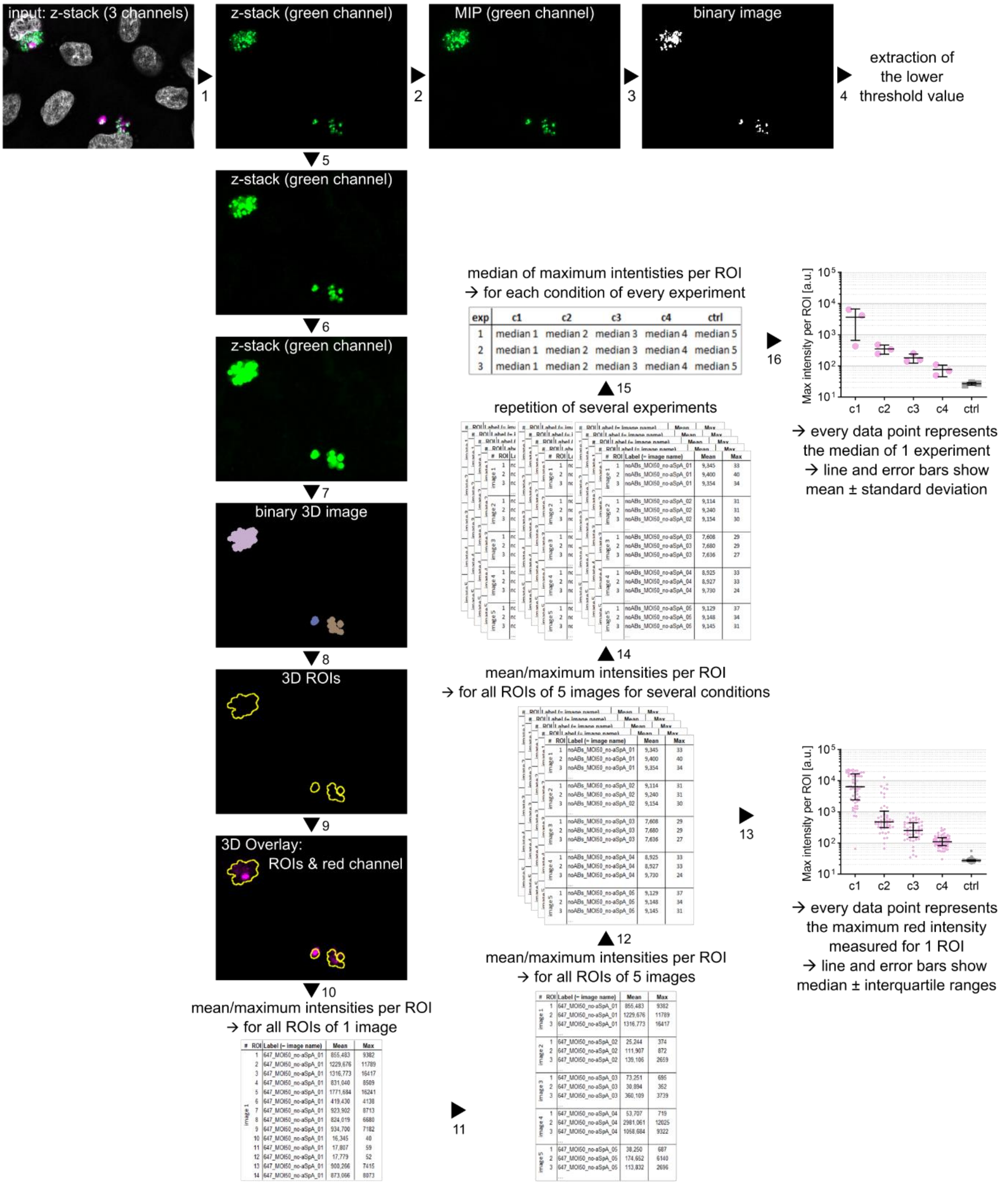


**Figure S4:** Image analysis pipeline for the quantification of nonspecific antibody staining of *S. aureus*-infected cells. 3-channel z-stacks were deconvolved with Zen blue. The resulting images were further analyzed using Fiji[1] as follows. 1: Extraction of the green channel. 2: Maximum intensity projection of the green channel. 3: Thresholding of the green channel MIP using Renyi's entropy. 4: Extraction of the lower threshold value for later 3D segmentation. 5: Processing of the original green z-stack with a 3D maximum filter. 6: Morphological 3D dilation[2]. 7: Segmentation of the green z-stack using the 3D ROI manager[3] with the in step 3 extracted lower threshold value and 65535 (maximum value for 16-bit images) as lower and upper borders for thresholding. 8: Extraction of ROIs from the binary 3D image. 9: Overlay of ROIs with the red channel. 10: Measurement of the mean and maximum red intensity within each ROI of the image. 11: Repetition of steps 1 to 10 for, in total, five images per condition. 12: Repetition of steps 1 to 11 for all conditions of an experiment. 13: Results of individual experiments: Plotting of maximum red intensities per ROI for five images per condition. 14: Repetition of steps 1 to

13 for several biological replicates. 15: Determination of the median values of each condition for every biological replicate. 16: Results of biological replicates: Plotting of these median values results in one data point per condition and biological replicate. a.u.: arbitrary units; c: condition; ctrl: control; exp: experiment; max: maximum; MIP: maximum intensity projection; ROIs: regions of interest.

**Figure S5**

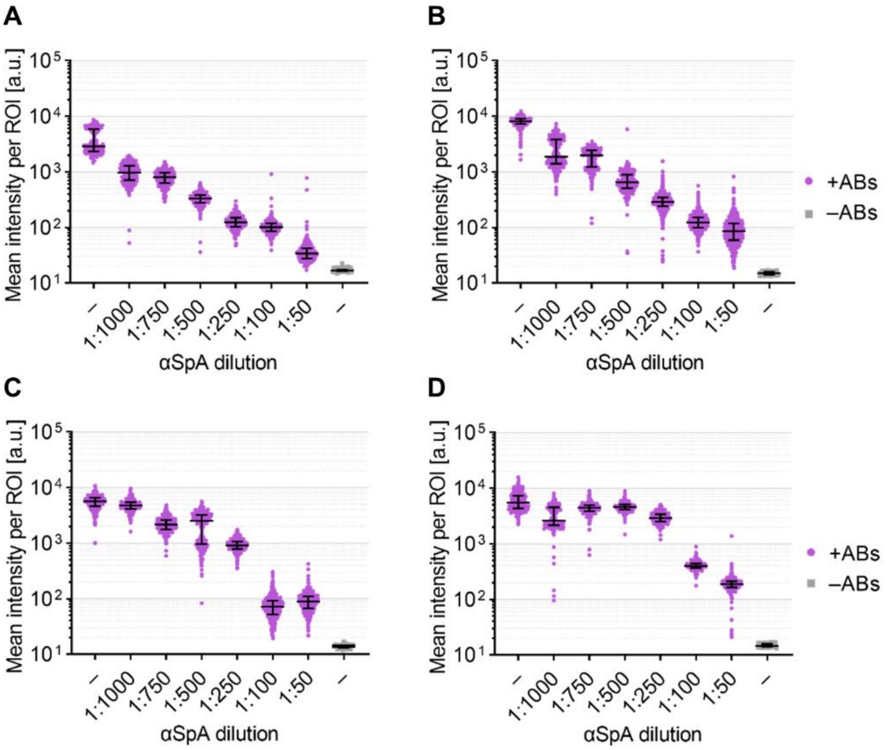


**Figure S5:** Pre-incubation with αSpA blocks nonspecific antibody staining of *S. aureus* – biological replicates underlying Figure 3B. (A-D) Four biological replicates of coverslip-coated GFP-expressing *S. aureus* were pre-incubated with different αSpA dilutions (1:1000 to 1:50) or with 3% BSA (–), and stained with the αNP/AF647 combination (+ABs), or without antibodies (–ABs). Quantification was done as described in Figure S2, steps 1-9. Five images per condition were analyzed. Line and error bars depict median ± interquartile range. a.u.: arbitrary units; ABs: antibodies; ROI: region of interest; SpA: *staphylococcal* protein A.

**Figure S6**

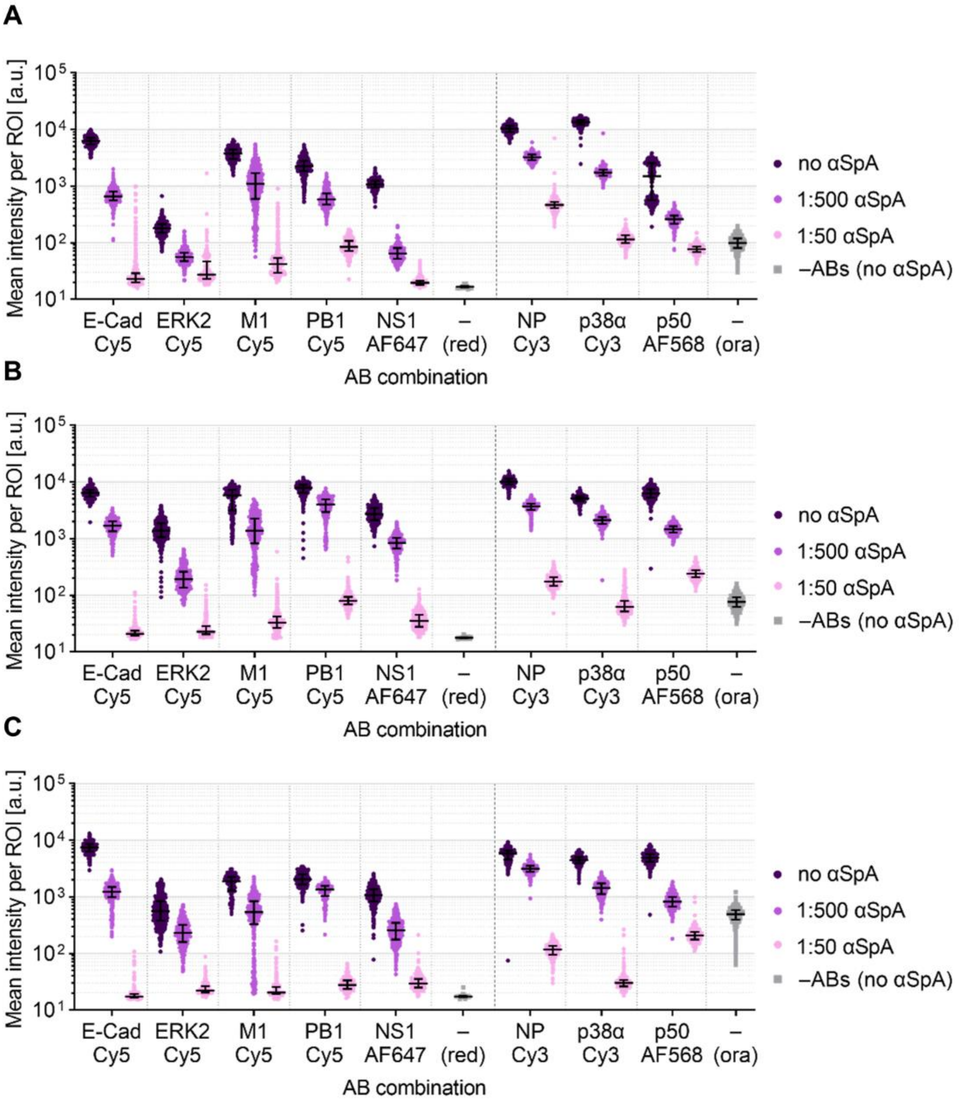


**Figure S6:** Pre-incubation with αSpA blocks nonspecific antibody staining of *S. aureus* – biological replicates underlying Figure 3C. (A-C) Three biological replicates of coverslip-coated GFP-expressing *S. aureus* were pre-incubated with αSpA or with BSA (no αSpA) and stained with different antibody combinations or without antibodies (–ABs). Quantification was done as described in Figure S2, steps 1-9. Five images per condition were analyzed. Line and error bars depict median ± interquartile range. a.u.: arbitrary units; ABs: antibodies; E-Cad: E-Cadherin; ERK2: extracellular signal-regulated kinase 2; M1: matrix protein 1; NP: nucleoprotein; NS1: non-structural protein 1; ora: orange; p38α: mitogen-activated protein kinase 14; p50: p50 subunit of NFκB; PB1: polymerase basic protein 1; ROI: region of interest; SpA: *staphylococcal* protein A.

**Figure S7**

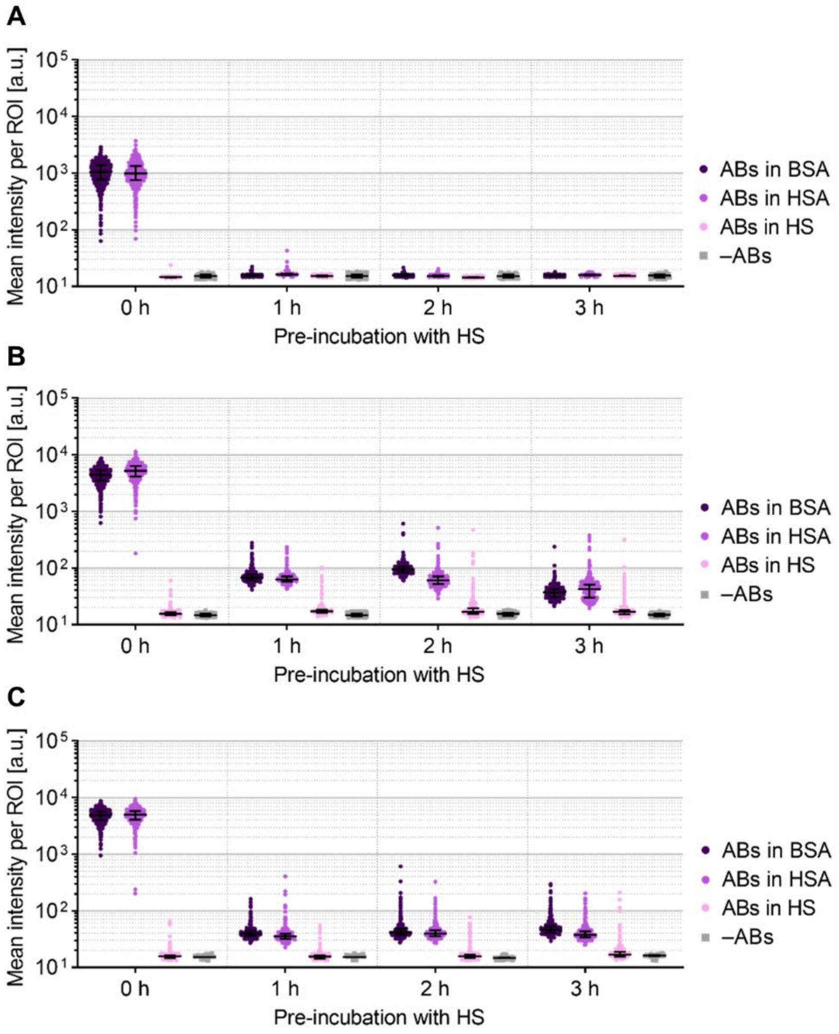


**Figure S7:** HS is a potent blocking solution and antibody diluent to prevent nonspecific antibody staining of *S. aureus* – biological replicates underlying Figure 4B. (A-C) Three biological replicates of coverslip-coated GFP-expressing *S. aureus* were pre-incubated with 100% HS for different incubation durations, and stained with the αNP/AF647 antibody combination. Antibodies were diluted in 3% BSA, 3% HSA, or 50% HS. No-antibody controls (–ABs) were pre-incubated with 3% BSA instead of HS. Quantification was done as described in Figure S2, steps 1-9. Five images per condition were analyzed. Line and error bars depict median ± interquartile range. a.u.: arbitrary units; ABs: antibodies; BSA: bovine serum albumin; HS: human serum; HSA: human serum albumin; ROI: region of interest.

**Figure S8**

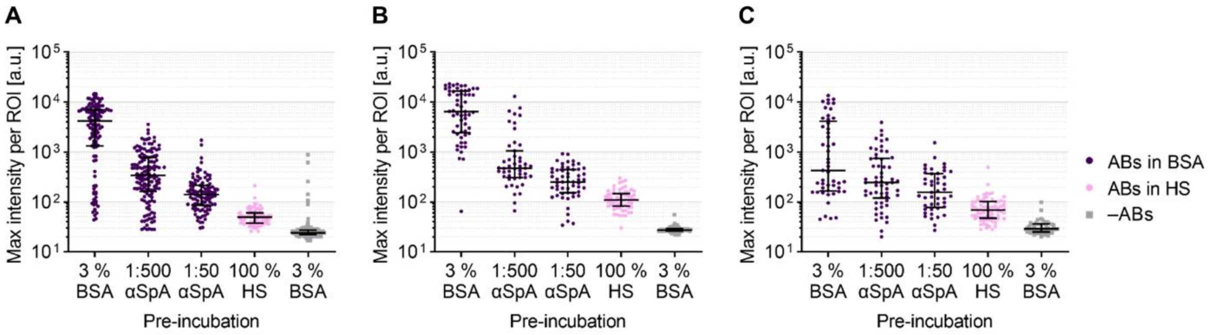


**Figure S8:** Validation of blocking strategies in *S. aureus*-infected cells – biological replicates underlying Figure 5B. (A-C) Three biological replicates of A549 cells were infected with GFP-expressing *S. aureus* and pre-incubated with BSA, αSpA, or HS. Cells were stained with the αNP/AF647 antibody combination and Hoechst 33342 diluted in 3% BSA or 50% HS, or only with Hoechst 33342 (–ABs). Quantification was done as described in Figure S4, steps 1-13. Five images per condition were analyzed. Line and error bars depict median ± interquartile range. a.u.: arbitrary units; ABs: antibodies; BSA: bovine serum albumin; HS: human serum; max: maximum; ROI: region of interest; SpA: *staphylococcal* protein A.

**Figure S9**

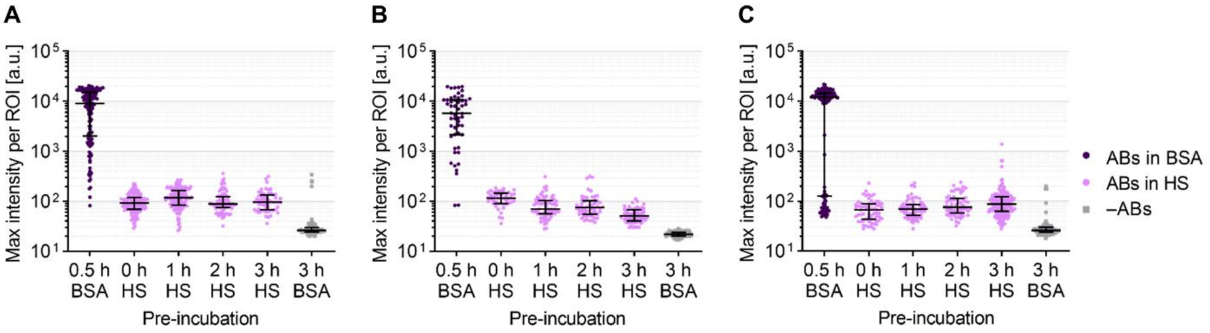


**Figure S9:** Validation of blocking strategies in *S. aureus*-infected cells – biological replicates underlying Figure 5C. (A-C) Three biological replicates of A549 cells were infected with GFP-expressing *S. aureus* and pre-incubated with 3% BSA or with 100% HS. Antibodies (αNP/AF647) and Hoechst 33342 were diluted in 3% BSA or 50% HS. No-antibody controls (–ABs) were only stained with Hoechst 33342. Quantification was done as described in Figure S4, steps 1-13. Five images per condition were analyzed. Line and error bars depict median ± interquartile range. a.u.: arbitrary units; ABs: antibodies; BSA: bovine serum albumin; HS: human serum; ROI: region of interest.
